# Hercules: a profile HMM-based hybrid error correction algorithm for long reads

**DOI:** 10.1101/233080

**Authors:** Can Firtina, Ziv Bar-Joseph, Can Alkan, A. Ercument Cicek

## Abstract

**Motivation:** Choosing whether to use second or third generation sequencing platforms can lead to trade-offs between accuracy and read length. Several studies require long and accurate reads including de novo assembly, fusion and structural variation detection. In such cases researchers often combine both technologies and the more erroneous long reads are corrected using the short reads. Current approaches rely on various graph based alignment techniques and do not take the error profile of the underlying technology into account. Memory- and time-efficient machine learning algorithms that address these shortcomings have the potential to achieve better and more accurate integration of these two technologies.

**Results:** We designed and developed *Hercules*, the first machine learning-based long read error correction algorithm. The algorithm models every long read as a profile Hidden Markov Model with respect to the underlying platform’s error profile. The algorithm learns a posterior transition/emission probability distribution for each long read and uses this to correct errors in these reads. Using datasets from two DNA-seq BAC clones (CH17-157L1 and CH17-227A2), and human brain cerebellum polyA RNA-seq, we show that Hercules-corrected reads have the highest mapping rate among all competing algorithms and highest accuracy when most of the basepairs of a long read are covered with short reads.

**Availability:** Hercules source code is available at https://github.com/BilkentCompGen/Hercules

## 1 1 Introduction

Next Generation Sequencing (NGS) technologies have revolutionized the field of genomics, and yet they suffer from two fundamental limitations. First and foremost, no platform is yet able generate a chromosome-long read. Average read length ranges from 100 bp to 10 Kbp depending on the platform. Second, reads are not error-free. The most ubiquitous platform, Illumina, produces the most accurate (<0.1% error rate), yet the shortest (100-150 bp) reads. Short read lengths present challenges in accurate and reproducible analyses [1, 2], as well as in building reliable assemblies [3–5]. On the other hand, Pacific Biosciences Single Molecule, Real-Time (SMRT) sequencing technology is capable of producing reads approximately 10 kbp-long on average with a substantially higher (~15%) error rate [6]. Similarly, Oxford Nanopore Technologies (ONT) platform can generate longer reads (up to ~900 Kbp) with the price of an even higher error rate (>20%) [7].

Complementary strengths and weaknesses of these platforms make it attractive to use them in harmony, i.e. merging the advantage of the longer reads generated by PacBio and ONT platforms with the accuracy of Illumina. Arguably, one can achieve high basepair accuracy using only PacBio or ONT reads, if the sequence coverage is sufficiently high [8]. However, the relatively higher costs associated with long read platforms make this approach prohibitive. Therefore, it is still more feasible to correct the errors in long reads using more accurate short reads in a hybrid manner.

There are two major hybrid approaches for long read error correction. First approach starts with aligning short reads onto long reads generated from the same sample, implemented by several tools such as PacBioToCA [9], LSC [10], proovread [11], and Colormap [12]. Leveraging the relatively higher coverage and accuracy of short reads, these algorithms fix the errors in long reads by calculating a consensus of the short reads over the same segment of the nucleic acid sequence. The second approach aligns long reads over a de Bruijn graph constructed using short reads, and the k-mers on the de Bruijn graph that are connected with a long read are then merged into a new, corrected form of the long read. Examples of this approach are LoRDEC [13], Jabba [14], HALC [15], and LoRMA [16].

The alignment based approach is highly dependent on the performance of the aligner. Therefore, accuracy, running time, and memory usage of an aligner will directly affect the performance of the downstream correction tool. The use of de Bruijn graphs eliminates the dependence on external aligners, and implicitly moves the consensus calculation step into the graph construction. However, even with the very low error rate in short reads and the use of shorter k-mers when building the graph, the resulting de Bruijn graph may contain bulges and tips, that are typically treated as errors and removed [17]. Accurate removal of such graph elements depend on the availability of high coverage data to be able to confidently discriminate real edges from the ones caused by erroneous k-mers [18, 19]. Therefore, performance of de Bruijn graph based methods depends on high sequence coverage.

Here we introduce a new alignment-based hybrid error correction algorithm, *Hercules*, to improve basepair accuracy of long reads. Hercules is the first machine learning-based long read error correction algorithm. The main advantage of Hercules over other alignment-based tools is the novel use of profile hidden Markov models (dubbed profile HMM or pHMM). Hercules models each long and erroneous read as a template profile HMM. It uses short reads as observations to train the model via Forward-Backward algorithm [20], and learns posterior transition and emission probabilities. Finally, Hercules decodes the most probable sequence for each profile HMM using Viterbi algorithm [21]. Profile HMMs are first popularized by the HMMER alignment tool for protein families [22] and are used to calculate the likelihood of a protein being a member of a family. To the best of our knowledge, this is the first use of a profile HMM as template with the goal of learning the posterior probability distributions.

All alignment-based methods in the literature, (i) are dependent on the aligner’s full CIGAR string for each basepair to correct, (ii) have to perform majority voting to resolve discrepancies among short reads, and (iii) assume all error types (i.e., deletion) are equally likely to happen. Whereas, Hercules only takes the starting positions of the aligned short reads from the aligner, which minimizes the dependency on the aligner. Then, starting from that position, it sequentially and probabilistically accounts for the evidence provided per short read, instead of just independently using majority voting per base-pair. Finally, prior probabilities for error types can be configured based on the error profile of the platform to be processed. As prior probabilities are not uniform, the algorithm is better positioned to predict the posterior outcome. Thus, it can also distinguish long read technologies.

We tested Hercules on the following datasets: (i) two BAC clones of complex regions of human chromosome 17, namely, CH17-157L1 and CH17-227A2 [6], and (ii) human brain cerebellum polyA RNA-seq data that LSC provides [10]. As the ground truth, (i) for BAC clones, we used finished assemblies generated from Sanger sequencing data from the same samples and (ii) for human brain data we used GRCh38 assembly. We show that Hercules-corrected reads has the highest mapping rate among all competing algorithms. We report ~20% improvement over the closest de Bruijn based method, ~2% improvement over the closest alignment-based method, and ~30% improvement over the original version of long reads. We also show that, should long reads have high short read breadth of coverage provided by the aligner (i.e., 90%), Hercules outputs the largest set of most accurate reads (i.e., ≥95% accuracy).

## 2 Methods

### 2.1 Overview of Hercules

Hercules corrects errors (insertions, deletions, and substitutions) present in long read sequencing platforms such as PacBio SMRT [23] and Oxford Nanopore Technologies [7], using reads from a more accurate orthogonal sequencing platform, such as Illumina [24]. We refer to reads from the former are referred as “long reads” and the latter as “short reads” in the remainder of the paper. The algorithm starts with preprocessing the data and obtains the short-to-long read alignment. Then, for each long read, Hercules constructs a pHMM template using the error profile of the underlying platform as priors. It then uses the Forward-Backward algorithm to learn the posterior transition/emission probabilities, and finally, uses the Viterbi algorithm to decode the pHMM to output the corrected long read (Figure 1).

**Figure 1:**
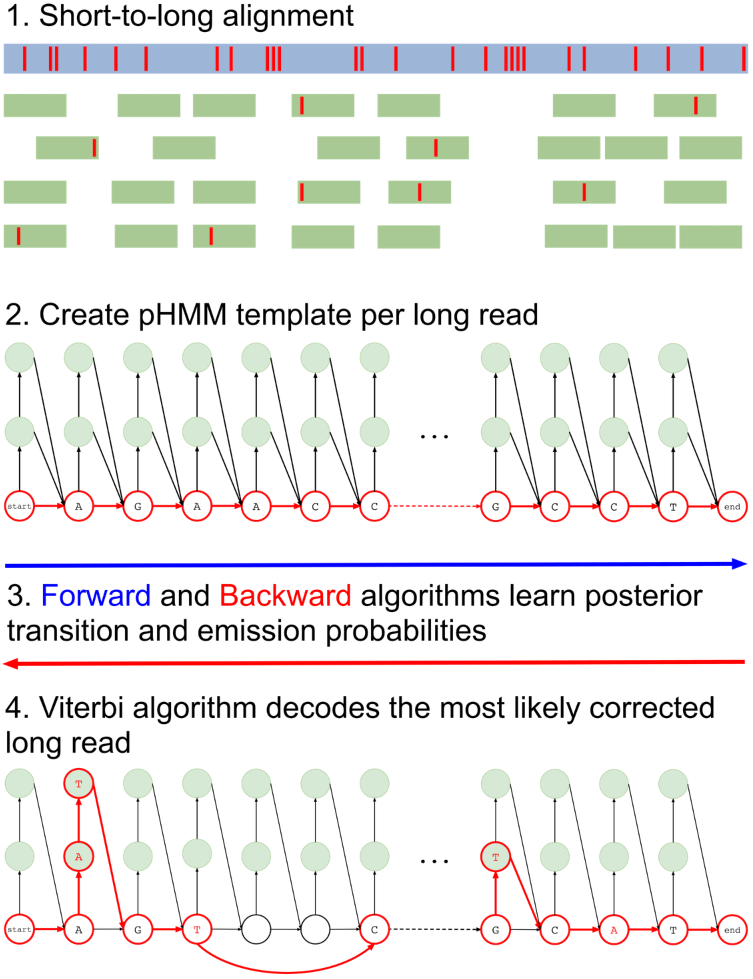
Overview of the Hercules algorithm. At the first step (1), short reads are aligned to long reads using an external tool. Here, red bars on the reads correspond to erroneous locations. Then, for each long read Hercules creates a pHMM with priors set according to the underlying technology as shown in (2). Using the starting positions of the aligned short reads, Forward-Backward algorithm learns posterior transition and emission probabilities. Finally, Viterbi algorithm finds the most likely path of transitions and emissions as highlighted with red colors in (3). The prefix and the suffix of the input long read in this example is “AGAACC…GCCT”. After correction, substring “AT” inserted right after the first “A”. Third “A” is changed to “T” and following two base-pairs are deleted. Note that deletion transitions are omitted other than this arrow, and only two insertion states are shown for clarity of the figure. On the suffix, a “T” is inserted and second to last base-pair is changed from “C” to “A”.

### 2.2 Preprocessing

#### 2.2.1 Compression

Similar to the approach of LSC [10], Hercules starts with compressing both short and long reads using a run-length encoding scheme. That is, it compresses repeating base-pairs to a single base-pair (e.g., AAACTGGGAC → ACTGAC). This is because PacBio and ONT platforms are known to produce erroneous homopolymer runs, i.e. consecutive bases may be erroneously repeated [7, 23], which affects the short read alignment performance drastically. Hercules, then, recalculates the original positions based on the original long reads. Even though this option is performed by default, it is still possible to skip the compression phase. Hercules obtains 3-fold increase in the most accurate reads (i.e., >95% accuracy) when compression phase is enabled. Doing so, the number of aligned reads also increases by ~30%.

#### 2.2.2 Short Read Filtering

After the step described above, the nominal length of some compressed reads may be substantially shortened. This would in turn cause such reads to be ambiguously aligned to the long reads, which is already a common problem in short read sequencing due to repeats [1, 2]. To reduce this burden, Hercules removes any compressed short read, if its number of non-N characters is less than a specified threshold (set to 40 by default). In addition to PCR and optical duplicates inherent in short read data, it is also highly likely that compression step will generate new duplicate sequences. These duplicate reads will not carry any extra information for correcting the long reads. Therefore, after compression, Hercules may remove these duplicates using a Bloom filter with 0.0001 false positive rate, keeping only a single read as the representative of the group. Duplicate removal is an optional phase, and we do not remove duplicates by default.

#### 2.2.3 Alignment and Decompression

Hercules is an alignment-based error correction tool. It outsources the alignment process to a third party aligner. It is compatible with any aligner that provides output in SAM format [25]. However, we suggest mappers that can report multiple possible alignments such as Bowtie2 [26] as we would prefer to cover most of the long read with short reads. Even though this might cause incorrect mappings, Hercules is able to probabilistically downplay the importance of such reads during the learning stage given sufficient short read depth. Hercules also assumes that the resulting alignment file is sorted. Thus, the file in SAM/BAM format must be coordinate sorted in order to ensure a proper correction with Hercules. Alignments are calculated using either compressed or original long and short reads (those that pass filtering step for the latter) as described in previous sections. After receiving the alignment positions from the aligner, Hercules decompresses both short and long reads and recalculates alignment start locations.

All other alignment-based error correction methods in the literature use the CIGAR string provided by the aligner. CIGAR string specifies where each basepair of the short read is mapped on the long read and where insertions and deletions have occurred. Then, per base-pair on the long read, they perform a majority voting among covering short reads. Hence, *learning* of correct basepairs is actually performed by the aligner, which makes the performance of such tools fully dependent on the aligner choice. Even though, Hercules is also an alignment-based method, the only information it receives from the aligner is the starting position of the mapping. Forward algorithm learns posterior probabilities starting from that position and it can go beyond the covered region. Thus, despite using the starting point information from the aligner, the algorithm can decide the short read is aligned after that point and also that the alignment is essentially different than what is claimed by the aligner in the CIGAR string. This minimizes the dependency of the algorithm on the aligner and makes Hercules the first alignment-based approach to reclaim the consensus learning procedure from the aligner.

### 2.3 The Profile HMM Structure

Hercules models each long read as a pHMM. In a traditional profile HMM [22], there are three types of states: deletion, insertion and match (mismatch) states. The aim is to represent a family of proteins (amino acid sequences) and then to use it to decide if an unknown protein is likely to be generated from this model (family). Our goal in representing a long read as a *template* pHMM is entirely different. For the first time, we use short reads that we know are generated from the source (e.g., same section of RNA/DNA), to update the model (not the topology but the transition and emission probabilities). So, the goal in the standard application is to calculate the likelihood of a *given* sequence, whereas in our application there is no given sequence. We would like to calculate the most likely sequence the model would produce, among all possible sequences that can be produced using the letters in the alphabet Σ. This requires us to impose some restrictions on the standard pHMM concept. First, we remove the self loop over insertion states. Instead, our model has multiple insertions states per position (basepair) on the long read. The number of insertion states is an input the algorithm. Second, we substitute deletion states with *deletion transitions*. The number of possible consecutive basepairs that can be deleted is an input parameter to the algorithm as well.

After we construct the model, we use the error profile of the underlying technology to initialize the prior transition and emission probabilities. Hercules first learns posterior transition and emission probabilities of each “long read profile HMM” using short reads that align to the corresponding long read. Then, it decodes the pHMM using the Viterbi algorithm [21] to generate the most probable (i.e. corrected, or consensus) version of the long read.

It should be noted that the meanings of insertion and deletion is reversed in the context of the pHMM. That is, an insertion state would insert a basepair into the erroneous long read. However, this means that the original long read did not have that nucleotide and had a deletion in that position. Same principle also applies to deletions.

Next, we first define the structure of the model (states and transitions) and explain how we handle different types of errors (i.e. substitutions, deletions, and insertions). We, then, describe the training process. Finally, we explain how Hercules decodes the model and generates the consensus sequence.

#### 2.3.1 Match States

Similar to traditional pHMM, we represent each basepair in the long read by a *match* state. There are four emission possibilities (Σ = {*A, C, G, T*}). The basepair that is observed in the uncorrected long read for that position *t* is initialized with the highest emission probability (*β*), while the probabilities for emitting the other three basepairs are set to the expected substitution error rate for the long read sequencing technology (*δ*) (Note that *β* >>*δ* and *β* + 3*δ* = 1).

We then set transition probabilities between consecutive match states *t* and *t* + 1 as “match transitions” with probability *α_M_*. Figure 2 exemplifies a small portion of the profile HMM that shows only the match states and match transitions. From a match state at position *t*, there are also transitions to (i) the first insertion state at position 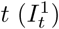, and (ii) to all match states at positions *t* + 1 + *x* where 1 ≤ *x* ≤ *k* and *k* determines the number of possible deletions.

**Figure 2:**
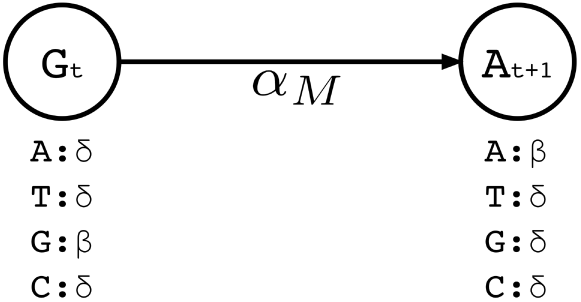
A small portion of the profile HMM built by Hercules. Here, two match states are shown, where the corresponding long read includes *G* at location *t*, followed by nucleotide *A* at location *t* + 1. At state *t*, the emission probability for *G* is set to *β*, and emission probabilities for *A*, *C*, *T* are each set to the substitution error rate *δ*. Match transition probability between states *t* and *t* + 1 is initialized to *α*_*M*_.

#### 2.3.2 Insertion States

Insertion states have self-loop transitions in standard profile HMMs to allow for multiple insertions at the same site, which creates ambiguity for error correction for two reasons. First, self-loops do not explicitly specify the maximum number of insertions. Thus, it is not possible predetermine how many times that a particular self-loop will be followed while decoding. Second, each iteration over the loop has to emit the same basepair. Since decoding phase prefers most probable basepair at each state, it is not possible for a standard profile HMM to choose different nucleotides from the same insertion state.

Instead of a single insertion state with a self loop per basepair in the long read, we construct *l* multiple insertion states for every match state at position *t* (e.g., 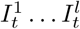). We replace self-loops with transitions between consecutive insertion states 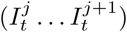 for position *t* (see Figure 3). The probability for each of such transitions is α_I_ and the number of insertion states per position *l* determines the maximum number of allowed insertions between two match states, which are set through user-specified parameters. All insertion states at position *t* have transitions to the match state at positions *t* + 1 (end states is also considered as a match state). For 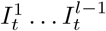 the probability of those transitions are α_*M*_, and for 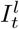, it is *α*_*M*_ + *α*_*I*_. All those states also have transitions to all match states at positions *t* + 1 + *x* where 1 ≤ *x* ≤ *k* and *k* determines the number of possible deletions.

**Figure 3:**
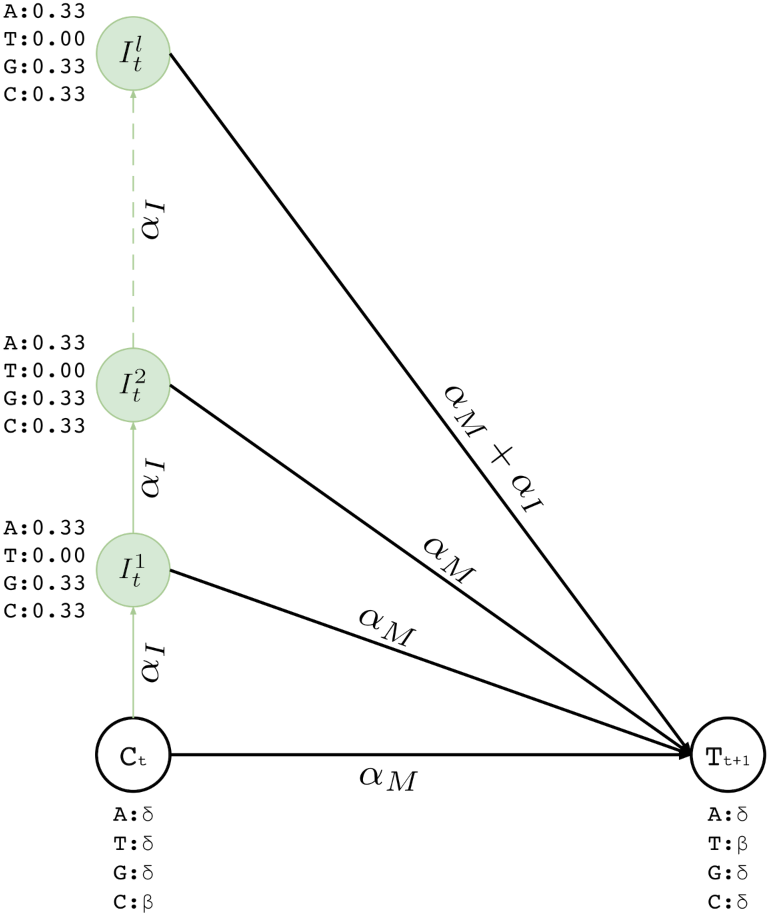
Insertion states for position *t*. Here we show two match states (*C_t_* and *T_t+1_*) and *l* insertion states 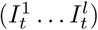. The number of insertion states limit the insertion length to at most *l* after basepair *t* of the corresponding long read. We also incorporate equal emission probabilities for all basepairs, except for the basepair represented by the corresponding match state *t* + 1 (*T* in this example).

We also set emission probabilities for the insertion states and assume they are equally likely. However, for the insertion states at position *t*, we set the probability of emitting the nucleotide *X* ∊ Σ to zero, if *X* is the most likely nucleotide at the match state of position *t* + 1. This makes the insertion more likely to happen at the last basepair of the homopolymer run. Otherwise, the likelihood of inserting a basepair is shared among all insertions states of the run, and it is less likely for any them to be selected during the decoding phase. All other basepairs have their emission probabilities set to 0.33 (i.e. 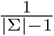) - see Figure 3.

#### 2.3.3 Deletion Transitions

In standard pHMM, a deletion state needs to be visited to skip (delete) a basepair, which does do not emit any characters. However, as described in the following sections, calculation of the forward and backward probabilities for each state is based on the basepair of the short read considered at that time. This means at each step, the Forward-Backward algorithm *consumes* a basepair of the short read to account for the evidence the short read provides for that state. As deletion states do not consume any basepairs of the short read, this in turn results in an inflation on the number of possibilities to consider and substantially increases the computation time. In an extreme case, the short read implying its mapped region on the long read may be deleted entirely. Since we are using an external aligner, such extreme cases are unlikely. Thus, we model deletions as transitions, instead of having deletion states. In our model, a transitions to (1 + *x*)-step away match states is established to delete *x* basepairs. As shown in Figure 4, match and insertion states at *t^th^* position have a transition to all match states at positions *t* + 1 + *x*, where 1 ≤ *x* ≤ *k* and *k* is an input parameter determining maximum number of deletions per transition. We calculate the probability of a deletion of *x* basepairs, 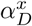, as shown in Equation 1.

**Figure 4:**
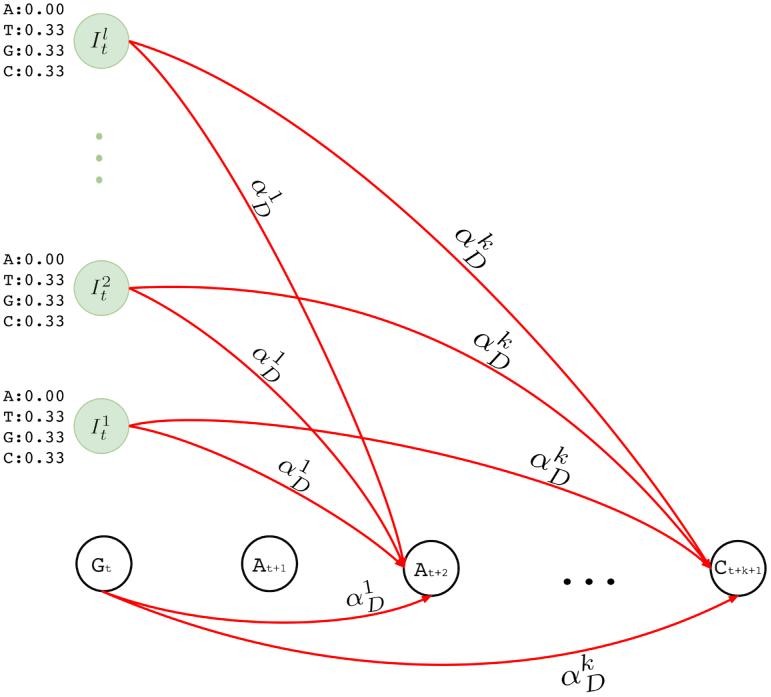
Deletion transitions (red) in Hercules pHMM. Insertion states of position *t* have the same deletion transitions with the match state of the same position, *t*. Any transition from position *t* to *t* + 1 + *x* removes *x* characters, skipping the match states between *t* and *t* + 1 + *x*, with 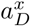 probability, where 1 ≤ *x* ≤ *k*

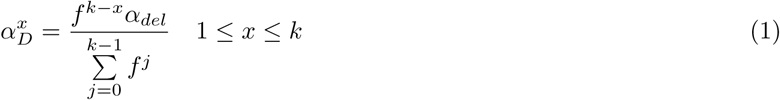

Equation 1 is a normalized version of a polynomial distribution where *α*_*del*_ is the overall deletion probability (i.e. *α*_*del*_ = 1 − *α*_*M*_ — α_*I*_), and *f* ∊ [1, ∞). As *f* value increases, probabilities of further deletions decrease accordingly.

#### 2.3.4 Training

An overall illustration of a complete pHMM for a long read of length *n* is shown in Figure 5 where (i) match states are labeled with *M_t_* where *M* ∊ Σ, (ii) insertion states are labeled with and 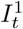, and 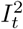 (iii) deletion transitions for *k* = 1 are shown. Note that *t^th^* basepair of a long read has one match state and two insertion states where 1 ≤ *t* ≤ *n*. There are also deletion transitions from every state at *t^th^* position to (*t* + 2)^*th*^ match state where (*t* + 2) ≤ *n*. This example structure allows only one character deletion at one transition since it is only capable of skipping one match state.

**Figure 5:**
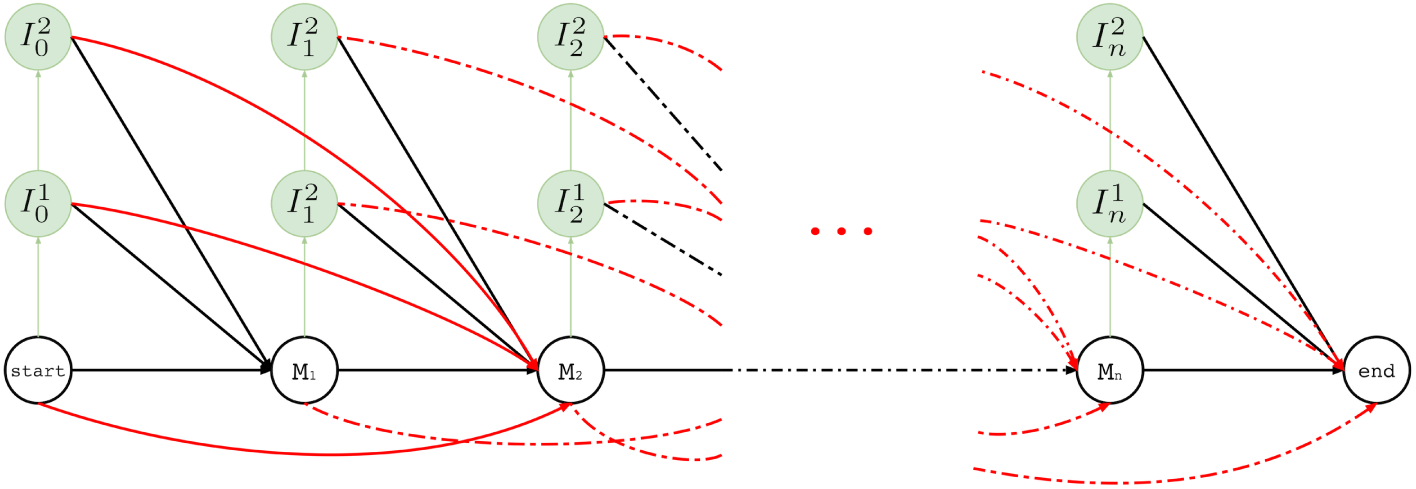
Hercules profile HMM in full. Here we show the overall look at the complete graph that might be produced for a long read *M* where its *t^th^* character is *M_t_* and *M_t_* ∊ {*A, T, G, C*} 1 ≤ *t* ≤ *n* and *n* is the length of the long read M (i.e. |*M*| = *n*). In this example, there are *n* many match states and two insertion states for each match state. Only one character deletion is allowed at one transition because transitions may only skip one match state.

Let the complete pHMM for a long read be the graph *G*(*V, E*). Per each mapped short read *s*, we extract a sub-graph *G_s_* (*V_s_*, *E_s_*). Here *G_s_* corresponds to the covered region of the long read with that short read. We train each *G_s_* using the Forward-Backward algorithm [20]. Hercules may not consider some of the short reads that align to same position in a long read to reduce computational burden. We provide *max coverage*, *mc*, option to only consider mc many number of short reads for a position during a correction, where *mc* = 20 by default. Short reads are selected based on their edit distance.

States that will be included in *V_s_* are determined by several factors. First, assume that s is aligned to start from the *q^th^* character of the long read. First, all match and insertion states between positions [*q* − 1, *q* + *m*) are included, where *m* is the length of the short read. If there are insertion errors, deletion transitions might be followed which may require training phase to consider *r* more positions where *r* is a parameter to the algorithm, which is fixed to ⸢*m*/3⸣. Thus all match and insertion states between [*q* + *m*, *q* + *m* + *r*) are also added. Finally, the match state at position *q* + *m* + *r* is also included (end state), and *M*_*q*−1_ acts as the start state. Every transition *E_ij_* ∊ *E* connecting state *i* and state *j* are included in *E_s_* if *i* ∊ *V_s_* and *j* ∊ *V_s_*. Each *E_ij_* ∊ *E_s_* is associated with a transition probability *α*_*ij*_ as described in previous subsections. For every pair of states, *i* ∊ *V_s_* and *j* ∊ *V_s_*, *α*_*ij*_ = 0 if *E_ij_* ∉ *E_s_*.

Let *e_i_*(*s*[*t*]) be the emission probability of the *t^th^* basepair of *s* from state *i*. We recursively calculate forward probabilities *F_t_*(*i*), ∀*i* ∊ *V_s_*, where 1 ≤ *t* ≤ *m* and *F_1_* (*i*) is the base case.

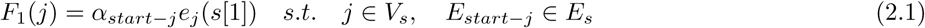

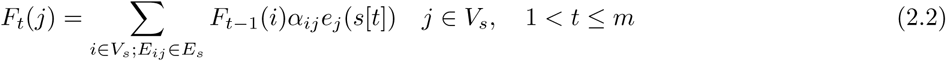

The forward probability *F_t_*(*j*) denotes the probability of being at state *j* having observed *t* basepairs of *s*. All transitions that lead to state *j* contributes to the probability with (*i*) the forward probability of the origin state *i* calculated with the *t* − 1^th^ basepair in *s*, multiplied by (ii) the probability of the transition from *i* to *j*, which is then multiplied by (iii) the probability of emitting *s*[*t*] from state *j*.

Backward probability *B_t_* (*j*) is also calculated recursively as follows, for each state *j* ∊ *V_s_* where *B_m_* (*j*) is the base case.

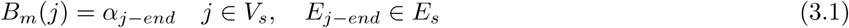

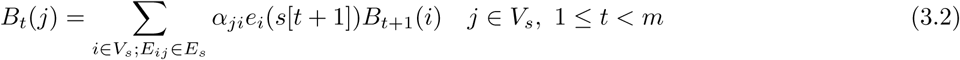

The backward probability *B_t_*(*i*) denotes the probability of being at state *j* and then observing the basepairs *t* + 1 to *m* from *s*. All outgoing transitions from state *j* to a destination state *i* contribute to this probability with (i) the backward probability of the state *i* with the *t* + 1^*th*^ basepair of *s*, multiplied by (ii) the transition probability from the state *j* to state *i*, which is then multiplied by (iii) the probability of *i* emitting the *t* + 1^*th*^ basepair of *s*.

During calculations of both forward and backward probabilities, at each time step *t*, we only consider *mf* number of highest forward probabilities of the previous time step, *t* − 1. This significantly reduces that time required to calculate forward and backward probabilities, while ensuring not to miss any important information if *mf* value is chosen as we suggest in the next section.

After calculation of the forward and backward probabilities (expectation step), posterior transmission and emission probabilities are calculated (maximization step), as shown in equations 4 and 5, respectively.

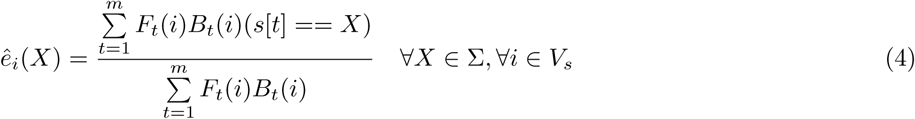

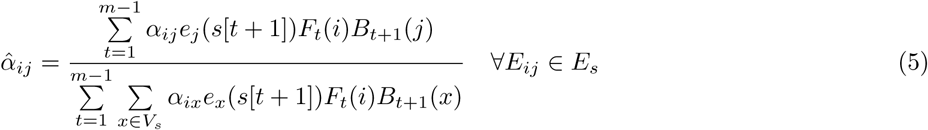

As we explain above, updates of emission and transition probabilities are exclusive for each sub-graph *G_s_*. Thus, there can be overlapping states and transitions that are updated within multiple sub-graphs, independently using the prior probabilities. For such cases, updated values are averaged with respect to posterior probabilities from each subgraph. Uncovered regions keep prior transition and emission probabilities, and *G* is updated with the posterior emission and transmission probabilities.

#### 2.3.5 Decoding

We use the Viterbi algorithm [21] to decode the consensus sequence. We define the consensus sequence as the most likely sequence the pHMM produces after learning new parameters. The algorithm takes *G* for each long read and finds a path from the start state to the end state, which yields the most likely transitions and emissions using a dynamic programming approach. For each state *j*, the algorithm calculates *υ_t_*(*j*), which is the maximum marginal forward probability *j* obtains from its predecessors when emitting the *t^th^* basepair of the corrected long read. It also keeps a back pointer, *b_t_*(*j*), which keeps track of the predecessor state *i* that yields the *v_t_*(*j*) value. Similar to calculations of forward and backward probabilities, we also consider *m f* many most likely Viterbi values from the previous time step, *t* − 1, to calculate *v*_*t*_(*j*).

Let *e_j_* ∊ Σ be the basepair that is most likely to be emitted from state *j* and *T* be the length of the decoded sequence which is initially unknown. The algorithm recursively calculates *v* values for each position *t* of a decoded sequence as described in equations 6.1 and 6.3. The algorithm stops at iteration *T*\* such that for the last *iter* iterations, the maximum value we have observed for *v*(*end*) cannot be improved - *iter* is a parameter and set to 100 by default. *T* is then set to *t*\* such that *v*_*t*\*_ (*end*) is the maximum among all iterations 1 ≤ *t* ≤ *T*\*.

1. Initialization

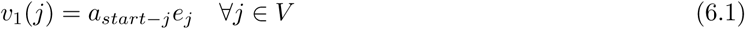

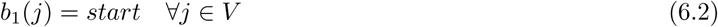
2. Recursion

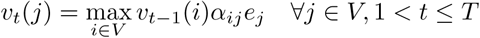

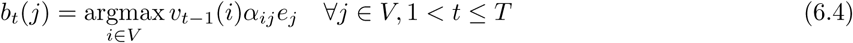
3. Termination

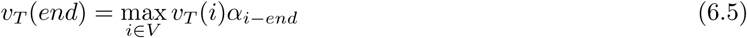

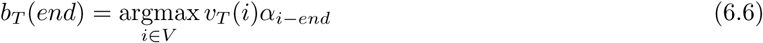

## 3 Results

### 3.1 Experimental setup

We implemented Hercules in C++ using the SeqAn library [28]. The source code is available under the BSD 3-clause license at https://github.com/BilkentCompGen/Hercules. We compare our method against Colormap, HALC, LoRDEC, LSC, and proovread. We ran all tools on a server with 4 CPUs with a total of 56 cores (Intel Xeon E7-4850 2.20GHz), and 1 TB of main memory. We assigned 60 threads to all programs including Hercules.

We used Bowtie2 with the multi-mapping option enabled to align the reads, and then we sorted resulting alignment files using SAMtools [25]. For Hercules, we set our parameters as follows: max insertion length (*l* = 3), max deletion length (*k* = 10), match transition probability (*a*_*M*_ = 0.75), insertion transition probability (*a*_*I*_ = 0.20), deletion transition probability (*a*_*del*_ = 0.05), deletion transition probability distribution factor (*f* = 2.5), match emission probability (*β* = 0.97), max coverage (*mc* =1), max filter size (*mf* = 100). We ran all other programs with their default settings.

In our benchmarks we did not include the error correction tools that use another correction tool internally such as LoRMA, which explicitly uses LoRDEC. Similarly, HALC also uses LoRDEC to refine its corrected reads. However, it is possible to turn off this option, therefore we benchmarked HALC without LoRDEC. We also exclude Jabba in our comparisons because it only provides corrected fragments of long reads and clips the uncorrected prefix and suffixes, which results in shorter reads. All other methods consider the entire long read. Additionally, several error correction tools such as LSC and proovread do not report such reads that they could not correct. Others, however, such as Hercules, LoRDEC, and Colormap report both corrected and uncorrected reads, preserving the original number of reads. In order to make the results of all correction tools comparable, we re-insert the original versions of the discarded reads to LSC and proovread output. We ran Colormap with its additional option (OEA) that incorporates more refinements on a corrected read. We observed that there was no difference between running OEA option or not in terms of the accuracy of its resulting reads, although Colormap with OEA option required substantially higher run time.

### 3.2 Data sets

To compare Hercules’s performance with other tools we used two DNA-seq and 1 RNA-seq data sets that were generated from two bacterial artificial chromosome (BAC) clones, previously sequenced to resolve complex regions in human chromosome 17 [6]. These clones are sequenced with two different HTS technologies: CH17-157L1 (231 Kbp) data set includes 93,785 PacBio long reads (avg 2,576bp; 1046X coverage) and 372,272 Illumina paired-end reads (76 bp each, 245X coverage); and CH17-227A2 (200 Kbp) data set includes 100,729 PacBio long reads (avg 2,663bp; 338X coverage) and 342,528 Illumina paired-end reads (76 bp each, 260X coverage). Additionally, sequences and finished assemblies generated with the Sanger platform are also available for the same BAC clones, which we use as the gold standard to test the correction accuracy. The RNA-seq data is generated from a human brain sample, which is also used in the original publication of the LSC algorithm [10]. This transcriptome data set includes 155,142 long reads (avg 879bp). The corresponding 64,313,204 short reads are obtained from the Human Body Map 2.0 project (SRA GSE30611). We calculated error correction accuracy using the human reference genome (GRCh38) for the human brain transcriptome data as the gold standard.

We report error-corrected PacBio reads generated from two BAC clones (CH17-157L1 and CH17-227A2) using Illumina reads from the same resource, and long reads from human brain transcriptome using Illumina reads from the Human Body Map 2.0 project. We report the accuracy as the alignment identity as calculated by the BLASR [27] aligner. *Mapped* refers to the number of any reads aligned to the Sanger-assembled reference for these clones with any identity, where the other columns show the number of alignments within respective identity brackets.

### 3.3 Correction accuracy

After correction, we align the corrected reads to the corresponding ground truth using BLASR [27] with the *noSplitSubreads* option, which forces entire read to align, if possible. Then, we simply calculate the accuracy of a read as reported by BLASR as alignment identity. We report the number of reads that align to the gold standard, and the alignment accuracy in four different accuracy brackets. We observe that the number of confident alignments were the highest in both BAC clones and the human brain transcriptome for the reads corrected by Hercules (Table 1). Furthermore, for CH17-157L1, Hercules returned the largest number of corrected reads with the highest (>95%) accuracy. In contrast, the LoRDEC had slightly more reads with the highest (>95%) accuracy for CH17-227A2 compared to Hercules. We further investigated the reasons for lower Hercules performance for this clone, and we found that only ~10% of the reads had > 90% short reads breadth coverage. For such reads with better coverage, Hercules’s performance surpassed that of LoRDEC and all other correction tools (Table 2).

**Table 1:** Summary of error correction

| Tool | Number of aligned reads |  |  |  |  |  |  |  |  |  |  |  |
| --- | --- | --- | --- | --- | --- | --- | --- | --- | --- | --- | --- | --- |
|  | CH17-157L1 BAC clone |  |  |  | CH17-227A2 BAC clone |  |  |  | Human brain transcriptome |  |  |  |
|  | Mapped | 80-90% | 90-95% | >95% | Mapped | 80-90% | 90-95% | >95% | Mapped | 80-90% | 90-95% | >95% |
| Uncorrected | 33,842 | 17,582 | 1,974 | 461 | 45,625 | 25,356 | 7,385 | 219 | 150,000 | 107,493 | 17,884 | 921 |
| Colormap | 36,391 | 13,586 | 7,904 | 6,235 | 46,209 | 18,398 | 12,364 | 4,715 | 150,663 | 49,803 | 35,412 | 50,518 |
| HALC | 35,220 | 17,867 | 2,974 | 2,249 | 46,680 | 23,969 | 9,662 | 2,356 | 150,970 | 93,500 | 29,822 | 6,682 |
| LoRDEC | 36,812 | 10,320 | 7,397 | 12,167 | 47,304 | 8,875 | 7,010 | **25,844** | 149,923 | 40,914 | 38,927 | 56,818 |
| LSC | 43,431 | 13,534 | 7,511 | 12,277 | 55,853 | 13,619 | 10,205 | 23,509 | 150,771 | 33,910 | 36,706 | **66,726** |
| Hercules | **44,229** | 13,609 | 7,569 | **12,583** | **56,140** | 13,433 | 9,809 | 24,269 | **151,903** | 34,360 | 38,352 | 65,880 |
| proovread | 38,962 | 14,918 | 6,714 | 7,956 | 47,344 | 17,460 | 9,233 | 11,140 | 150,235 | 82,344 | 21,891 | 25,944 |

**Table 2:** Correction accuracy given high breadth of coverage (> 90%) in CH17-227A2.

| Tool | No. of Aligned Reads |  |  |  |
| --- | --- | --- | --- | --- |
|  | Mapped | 80-90% | 90-95% | >95% |
| Uncorrected | 4,429 | 2,135 | 2,093 | 26 |
| Colormap | 4,432 | 668 | 2,345 | 1,271 |
| HALC | 4,433 | 1,789 | 2,436 | 58 |
| LoRDEC | 4,425 | 124 | 224 | 4,006 |
| LSC | 4,473 | 45 | 149 | 4,264 |
| Hercules | **4,476** | 43 | 142 | **4,273** |
| proovread | 4,434 | 1,722 | 1,665 | 880 |

### 3.4 Run time and memory requirements

Finally, we benchmarked computational requirements for each tool (Table 3). We observed that LoRDEC was the fastest algorithm, and the run times of alignment-based tools were similar. Running time of Hercules is still on the same scale even though the learning procedure is more time consuming because of per-read calls to Forward Backward and Viterbi algorithms. Since error correction is an offline and one-time task we argue that given the gain in accuracy this run time is acceptable.

**Table 3:**
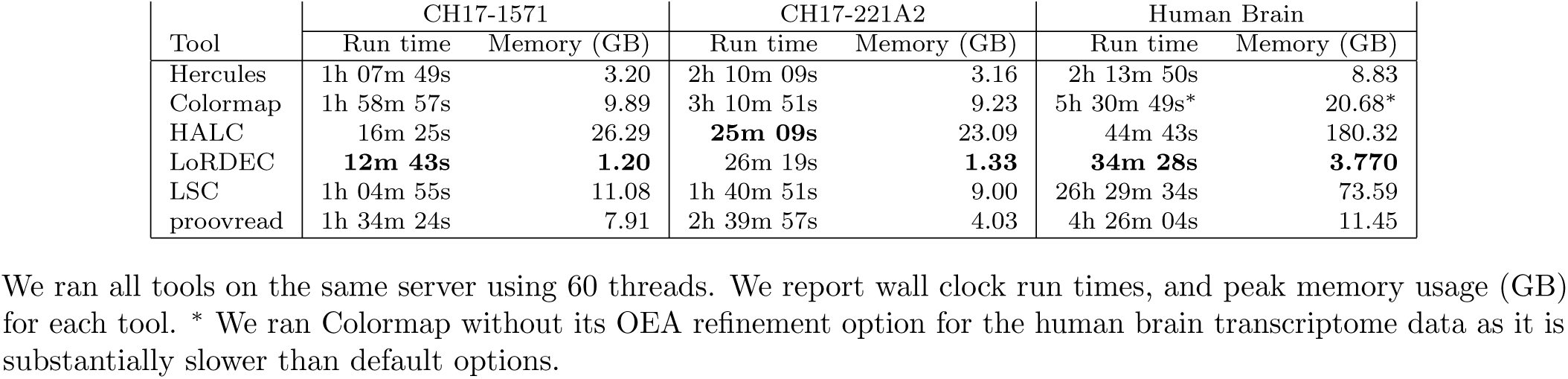
Requirements of computational resources for all methods we compared.

We also investigated the memory requirements of the tools we benchmarked (Table 3). We show that Hercules

The input included 9,449 PacBio reads which at least 90% of their length were covered by short reads. We report the accuracy as the alignment identity as calculated by the BLASR [27] aligner. *Mapped* column refers to the number of any reads aligned to the Sanger-assembled reference for this clone, where the other columns show the number of alignments within respective identity brackets. Uses only a modest amount of memory, second to only LoRDEC. In our experiment data sets, the maximum memory Hercules required was 8 GB, which makes it usable in commodity desktop computers. We note that the memory requirements of Hercules scale linearly with the lengths of the long reads to be corrected simultaneously in multiple threads, and the LoRDEC memory usage will depend on the size of the de Bruijn graph. Note that LoRDEC is alignment-free, and it builds a de Bruijn graph for the entire short read data set. Thus, its memory usage will be determined by the size of the input DNA (whole genome, whole transcriptome, or clones).

## 4 Discussion

Long reads such as PacBio are attractive alternatives to short Illumina reads due to the ability to span across common repeats, which present analytical and computational challenges to accurately characterize and assemble genomes. However, their error rate is also substantially higher than that of short reads, inferring additional challenges in basepair accuracy. Therefore, it is of utmost importance to correct short indel and substitution errors either using very high coverage long read data, which substantially increases sequencing cost, or orthogonal and less expensive short read sequencing data. Here we introduced a new algorithm, Hercules, as the first hidden Markov model based method to correct erroneous long reads using short but accurate Illumina data.

Profile HMMs were previously proven to be powerful for the problems in sequence analysis such as multiple sequence alignment and protein classification [29], however they were not exploited for error correction before. In this paper, we modified the standard pHMM to leverage their probabilistic basepair consensus representation to correct long reads given set of aligned short reads. Our proposed pHMM-like structure offers a flexible approach since its initial parameters can be redefined based on the error profile of any other error-prone sequencing technology, such as Oxford Nanopore. Moreover, although Hercules uses the short to long read information, it is largely independent of the underlying aligner, as it only uses the map start location, and uses the entire short read to train the pHMM.

Hercules is slower and more memory demanding compared to LoRDEC. However, as we have explained in the Results section, the memory requirement of LoRDEC will scale with the length of the input genome to be sequenced. In terms of run time, the main bottleneck for Hercules lies in the pHMM training for each long read. There are several algorithms proposed to improve running time and memory requirements of the standard Viterbi algorithm and Forward/Backward likelihood calculations that may be used to improve running time [30–32]. Additionally Hercules may further be accelerated through SIMD vectorization of pHMM training and decoding [33–35] that we leave as future work.

## Acknowledgements

We thank J. Huddleston for providing us with the BAC clone data sets, including their respective finished assemblies.

## Funding

Funding for this project was provided by a TÜBÍTAK grant (TÜBÍTAK-1001-215E172) and a Marie Curie Career Integration Grant (303772) to C.A., and a Turkish Academy of Sciences grant (TUBA-GEBIP) to A.E.C.

